# A landscape model for cell fate decisions during mesoendoderm differentiation in C. elegans based on Wnt dynamics

**DOI:** 10.1101/2021.06.09.447780

**Authors:** Shyr-Shea Chang, Zhirong Bao, Eric D. Siggia

## Abstract

Geometric models allow us to quantify topography of the Waddington landscape and gain quantitative insights of gene interaction in cell fate differentiation. Often mutant phenotypes show partial penetrance and there is a dearth of quantitative models that can exploit this data and make predictions about new allelic combinations with no additional parameters. *C. elegans* with its invariant cell lineages has been a key model system for discovering the genes controlling development. Here we focus on the differentiation of the endoderm founder cell named E from its mother, the EMS cell. Mutants that convert E to its sister MS fate have figured prominently in deciphering the Wnt pathway in worm. We construct a bi-valued Waddington landscape model that predicts the effect on POP-1/TCF and SYS-1/beta-catenin levels based on the penetrance of mutant alleles and RNAi, and relates the levels to fate choice decisions. A subset of the available data is used to fit the model and remaining data is then correctly predicted. Simple kinetic arguments show that contrary to current belief the ratio of these two proteins alone is not indicative of fate outcomes. Furthermore, double mutants within a background reduction of POP-1 levels are predicted with no adjustable parameters and their relative penetrance can differ from the same mutants with the wild-type POP-1 level, which calls for further experimental investigations. Our model refines the content of existing gene networks and invites extensions to other manifestations of the Wnt pathway in worm.

## Introduction

Development is a process of a single cell growing into a functional organism, and this requires a precise orchestration of cell fates (Wolpert and Tickle, 2011). Cell fates are known to be determined by various transcription factors and signaling events, and the underlying gene networks have been discovered in many systems (Zacharias and Murray, 2016; Katoh, 2007). While the structure of gene networks greatly enhanced our understanding on cell fate decisions, it only provides a qualitative description of the process. To improve our understanding, quantitative models with predictive power are much desired.

One such model can be obtained by fitting Waddington’s developmental landscape to experimental data (Waddington, 2014). This metaphor for development was recently given a mathematical formulation in (Rand et al., 2021; Corson et al., 2017). It abstracts from a gene network model the decision points and terminal states and represents the movements among them in geometric terms that are amenable to a compact parameterization. A landscape model was applied to vulva development in *C. elegans* and give quantitative predictions (Corson and Siggia, 2012; Corson and Siggia, 2017; Corson et al., 2017).

The mesoendoderm differentiation in the nematode *C. elegans* provides a great system to study cell fate decisions. Wild-type *C. elegans* embryos follow an invariant cell lineage (Sulston et al., 1983), which makes it an ideal model for observing cell fate differentiation. The mesoendoderm differentiation begins when the progenitor EMS cell divides into the mesoderm progenitor MS cell and the endoderm progenitor E cell. This process has been closely examined, with gene network largely uncovered and extensive data on fate differentiation under genetic mutations (Sawa and Korswagen, 2013; Maduro et al., 2007; Huang et al., 2007; Shin et al., 1999; Bei et al., 2002; Thorpe et al., 1997; Lin et al., 1995). The prior knowledge and experimental data enable a quantitative landscape model to be established for this developmental process.

Similar to its role in vertebrates, the Wnt pathway plays an essential role in the mesoendoderm differentiation in *C. elegans* (Sawa and Korswagen, 2013; Nusse and Clevers, 2017). The Wnt signal is transmitted to the posterior side of the EMS cell prior to cell division, activating MOM-5/Frizzled and DSH-2, MIG-5/Disheveled. This triggers WRM-1/beta-catenin and APR-1/APC to localize to the cortex in the anterior of the cell, which, along with other mechanisms, leads to asymmetric concentration of POP-1/TCF and SYS-1/beta-catenin in the MS and E cell nuclei. Quantitative live imaging to measure POP-1 and SYS-1 levels have been established (Zacharias et al., 2015), but systematic measurements across the genes, alleles and combination would be inefficient.

In this work we build a quantitative landscape model using Michaelis-Menten dynamics and train it with the experimental data. The model successfully fits the data on single and double mutant penetrance with a minimal number of parameters. Then we validate the model by verifying its predictions against testing data and observing the effect of perturbation on parameters. The model reveals quantitative aspects of the development, such as the nonlinear dependence of cell fate on the relative levels of POP-1 and SYS-1 and the changes in these levels caused by genetic mutations. These aspects lead to interesting predictions on fate differentiations in untested double mutants, especially when mutants have opposite effects on downstream transcription factors. Our work provides a first quantitative model for Wnt-induced cell fate decision in *C. elegans* with falsifiable predictions that can be experimentally tested.

## Results

In this work we focus on the fate differentiation of MS and E cells, the daughter cells of EMS cell (Fig. 1A). MS and E cells can assume a number of fates under different mutant backgrounds: the MS cell can adopt the E fate, or the fate of a cousin cell called C, and even a mixed fate between MS and C (Du et al., 2015; Owraghi et al., 2010), while the E cell can adopt the MS or C fate. A large amount of experimental data on fate differentiation focus on the fate outcomes of the E cell (Table S1), and specifically the E to MS transformation. and thus we build and train our model for this binary fate choice (Fig. 1B).

**Fig. 1:**
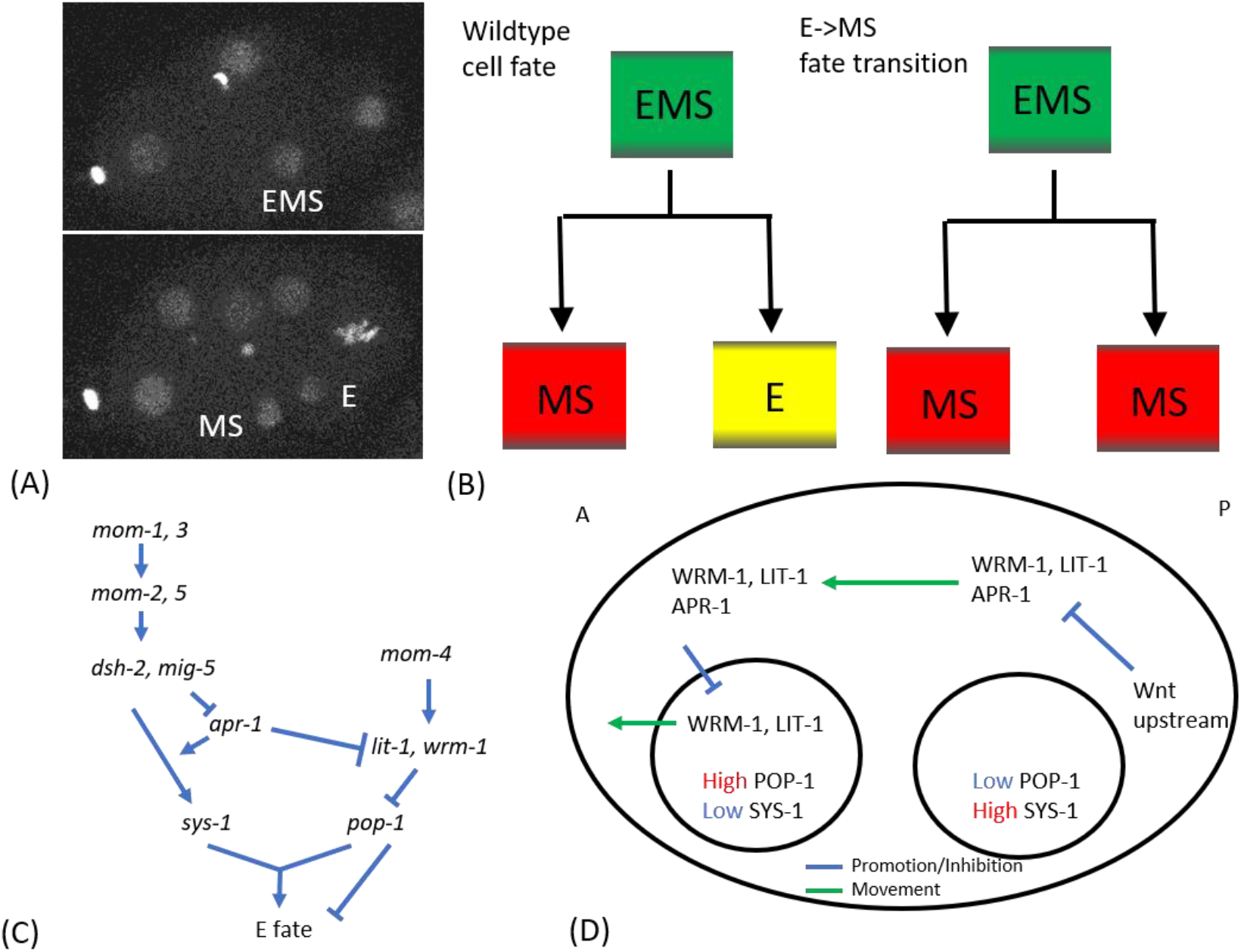
Overview on EMS differentiation and Wnt signaling. (A) Fluorescence images of EMS dividing into its daughter cells MS and E. (B) Cell lineage diagrams of wildtype cell lineage (left) and embryos with E->MS phenotype (right). (C) Gene network for Wnt signaling and E fate (adapted from Huang et al., 2007). (D) Spatial diagram of Wnt signaling and POP-1, SYS-1 levels in anterior and posterior nuclei.

The gene network determining the fate of E cell between E and MS fate is the Wnt signaling pathway (Fig. 1C). The downstream effectors are POP-1/TCF and SYS-1/beta-catenin, and the concentration ratio between these proteins is crucial for the fate outcome (Huang et al., 2007; Sawa and Korswagen, 2013). POP-1 and SYS-1 are regulated by different mechanisms. For POP-1 regulation, the posterior Wnt signal causes localization of WRM-1/beta-catenin and APR-1/APC to the anterior cortex, which reduces the amount of the WRM-1, LIT-1/Nemo complex from the anterior nucleus (Fig. 1D; Sawa, 2012). As the WRM-1, LIT-1 complex, facilitated by MOM-4/TAK1, phosphorylates POP-1 for exporting it from the nucleus, the down-regulation of nuclear WRM-1 and LIT-1 up-regulates nuclear POP-1 level, creating an asymmetry in POP-1. The mechanism for SYS-1 asymmetry is less known. It is believed that Wnt signal stabilizes SYS-1 in the posterior nucleus, protecting it from degradation (Sawa and Korswagen, 2013). From the Wnt signaling pathways we divide the genes into 3 groups in this work: Wnt upstream (*mom-1,2,3,5*), *sys-1* branch (*sys-1*), and *pop-1* branch (*mom-4, lit-1, wrm-1, pop-1*). This grouping comes from the topology of the gene network, as *mom-1,2,3,5* are upstream of all the Wnt signaling pathways, while *mom-4, lit-1, wrm-1* only regulates POP-1 level. Mutations in *pop-1* and *sys-1* also only affect POP-1 and SYS-1 respectively, and hence are included in their own branches. Combined together, these mechanisms lead to high POP-1 and low SYS-1 levels in the anterior nucleus, and low POP-1 and high SYS-1 levels in the posterior nucleus, which causes the anterior cell to admit MS fate and the posterior cell to admit E fate.

The data we use in this work is the quantitative data of penetrance on fate outcome for various single and double mutants. The penetrance is the percentage of cells under a certain genetic background admitting a fate differentiation, and we only consider E-to-MS fate differentiation here. Previous studies reported the penetrance of cell fate differentiation during EMS development under various mutations (Table 1; Bei et al., 2002, Huang et al., 2007; Shin et al., 1999; Thorpe et al., 1997; Maduro et al., 2005; Broitman-Maduro et al., 2009), and we aggregate these data in Table S1 for quick reference.

**Table 1:** Experimentally observed penetrance of the phenotype that E cell admits the fate of MS cell under genetic mutations. We highlight the mutants exhibiting “non-monotonicity” in red (see Results).

| Genotype | E->MS<br>penetrance<br>(%) | Reference | Training/Testin<br>g |
| --- | --- | --- | --- |
| Wildtype | 0 |  | Training |
| *sys-1(RNAi)* | 4 | Huang et al., 2007 | Training |
| *lit-1(t1534)* | 0 | Huang et al., 2007 | Training |
| *lit-1(t1512)* | 99 | Huang et al., 2007 | Training |
| *lit-1(RNAi)* | 96 | Shin et al., 1999 | Training |
| *wrm-1(RNAi)* | 100 | Huang et al., 2007 | Training |
| *mom-1(or10)* | 83 | Huang et al., 2007 | Training |
| *mom-2(or42)* | 72 | Huang et al., 2007 | Training |
| *mom-2(ne141)* | 53 | Shin et al., 1999 | Training |
| *mom-2(ne141 bei2002)* | 66 | Bei et al., 2002 | Training |
| *mom-2(RNAi)* | 28 | Huang et al., 2007 | Training |
| *mom-3(or78)* | 70 | Huang et al., 2007 | Training |
| *mom-4(ne19)* | 43 | Shin et al., 1999 | Training |
| *mom-4(or39)* | 40 | Thorpe et al., 1997 | Training |
| *mom-4(ne135)* | 11 | Shin et al., 1999 | Training |
| *mom-5(or57)* | 14 | Huang et al., 2007 | Training |
| *mom-5(zu193)* | 4 | Bei et al., 2002 | Training |
| *mom-5(RNAi)* | 2 | Shin et al., 1999 | Training |
| *apr-1(RNAi)* | 29 | Huang et al., 2007 | Training |
| *apr-1(RNAi shin1999)* | 42 | Shin et al., 1999 | Training |
| *dsh-2(RNAi);mig-5(RNAi)* | 4 | Bei et al., 2002 | Training |
| *mom-1(or10); sys-1(RNAi)* | 93 | Huang et al., 2007 | Training |
| *mom-2(or42); sys-1(RNAi)* | 100 | Huang et al., 2007 | Training |
| *mom-3(or78); sys-1(RNAi)* | 81 | Huang et al., 2007 | Training |
| *mom-4(ne19); sys-1(RNAi)* | 100 | Huang et al., 2007 | Training |
| *mom-5(or57); sys-1(RNAi)* | 60 | Huang et al., 2007 | Training |
| *lit-1(t1534); sys-1(RNAi)* | 100 | Huang et al., 2007 | Training |
| *wrm-1(RNAi); sys-1(RNAi)* | 100 | Huang et al., 2007 | Training |
| *mom-4(or39); mom-2(or42)* | 100 | Thorpe et al., 1997 | Testing |
| *mom-2(ne141);mom-4(ne19)* | 100 | Shin et al., 1999 | Testing |
| *mom-5(RNAi);mom-4(ne19)* | 100 | Shin et al., 1999 | Training |
| *mom-5(RNAi);mom-4(ne135)* | 100 | Shin et al., 1999 | Training |
| *lit-1(RNAi);mom-4(ne19)* | 95 | Shin et al., 1999 | Testing |
| *apr-1(RNAi shin1999) ;mom-4(ne19)* | 100 | Shin et al., 1999 | Testing |
| *dsh-2(RNAi);mig-5(RNAi); mom-2(ne141 bei2002)* | 7 | Bei et al., 2002 | Testing |
| *dsh-2(RNAi);mig-5(RNAi);mom-5(zu193)* | 8 | Bei et al., 2002 | Training |

### The landscape model

To model the fate differentiations during EMS development, we designed a landscape model inspired by the Waddington’s landscape metaphor for development (Waddington, 2014). The quantified landscape is represented by a curve in 1D space, where negative *x* indicates MS fate, positive *x* indicates E fate, and *x* close to zero indicates EMS fate. The landscape model has 2 key elements: (1) the transition from monostability to bistability following EMS division and (2) the linear bias toward MS or E fate from Wnt signaling.

In terms of the first element, the transition from monostability to bistability models the decay in SKN-1, a key transcription factor that induces and maintains EMS fate (Bowerman et al., 1992; Du et al., 2014). The experimentally observed decay of SKN-1 (Lin, 2003) is modeled by a gradual transition from one to two minima landscape (Fig. 2A), which opens up MS and E fates for both the MS and E cells.

**Fig. 2:**
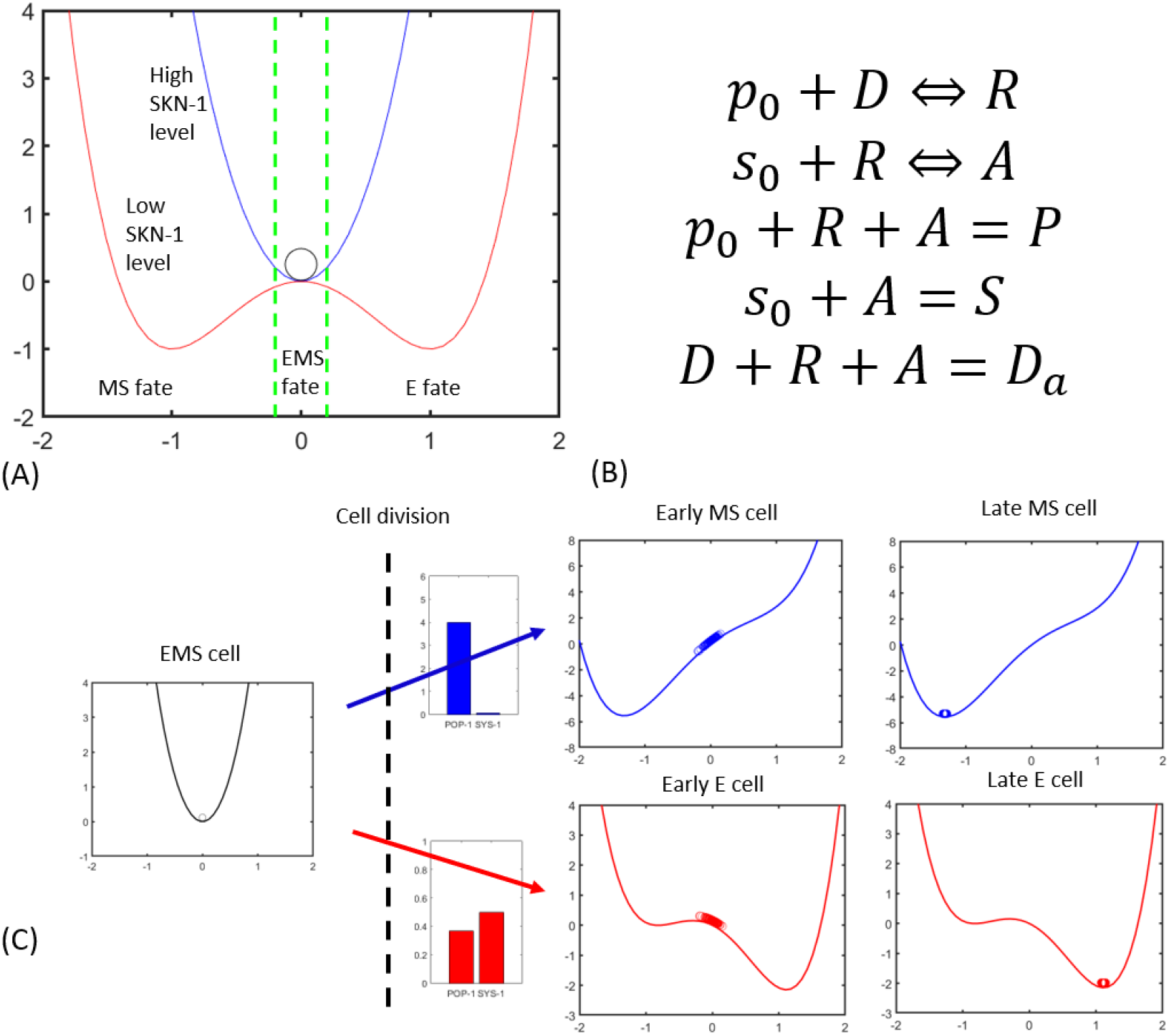
Overview on the landscape model with the Wnt dynamics. (A) The developmental landscape is quantified as a potential curve in the 1D space, with x=−1 and x=1 indicating MS and E fates. The MS and E fates are not available during the cell life of EMS (blue parabolic curve), and are opened up after EMS divides (red quartic curve). At the end of MS and E cell life, the fate of the cell is determined by its position (x<−0.2: MS fate. −0.2≤x≤0.2: EMS fate. x>0.2: E fate). The cell fate evolved downhill in the potential with additive noise to simulate partial penetrance. (B) The effect of Wnt signals is quantified by the Michaelis-Menten dynamics that link the POP-1 (P) and SYS-1 (S) levels to the active (A) and repressive (R) forms of the promoters of genes downstream of Wnt signaling. The other variables are *D*: unbound promoters, *p*_0_: unbound POP-1, *s*_0_: unbound SYS-1, and *D*_*a*_: the total amount of promoters. (C) The landscape model with the Wnt dynamics can then output cell fates from POP-1 and SYS-1 levels. The polynomial potentials are tilted by linear potentials determined by A, P from the Michaelis-Menten dynamics. For example, in wildtype MS cell, high POP-1 and low SYS-1 levels lead to a leftward bias toward MS fate (up), while in E cell, low POP-1 and high SYS-1 levels lead to a rightward bias toward E fate (down). The model can output penetrance of fate differentiations by simulating multiple cells at once with diffusion in cell fates.

The second element is designed to model the asymmetry between MS and E fate caused by Wnt signaling. To model this process, we explicitly write down the binding of Wnt downstream transcription factors, POP-1 and SYS-1, with the promoters of genes with major influence on fate outcomes, such as end-1, 3 (Maduro 2007 Dev. Bio.), by Michaelis-Menten kinetics (Fig. 2B). There are 2 chemical reaction equations and 3 equations for conservation laws. The first chemical reaction equation describes the binding of POP-1 to unoccupied promoters, resulting in the repressive form of the promoter. The second equation describes the binding of SYS-1 to the repressive form of the promoter, changing it to active. There is only one concentration variable per cell (no distinction between nucleus and cytoplasm) and we impose the conservation of POP-1, SYS-1 in each cell.

Given the levels of POP-1 and SYS-1, our model then calculates a linear potential based on the levels of active and repressive promoters, and adds it to the quartic potential after EMS division. An additive Gaussian noise in the equation describing relaxation in the potential completes the model and generates variable outcomes (see Methods, Eqn. (6)).

The model then includes these two elements to calculate the fate penetrance. The fate outcome is determined by the final position of the cell after it follows the potential gradient from the beginning of the EMS cell life to the end of MS and E cell life. For example, in the MS cell the POP-1 level is high while the SYS-1 level is low, resulting in a high level of repressive and a low level of activated promoters and hence a leftward-tilted potential (Fig. 2C, top). The fate of the EMS cell starts at *x* = 0, and is constrained to the EMS fate by the quadratic potential. After the potential becomes bistable and tilts leftward, the cell fate travels to the left following the potential, and eventually assumes MS fate. A similar scenario plays out for the E cell, except that high SYS-1 and low POP-1 levels lead to a rightward-tilted potential (Fig. 2C, bottom). In each run of the landscape model, we simulate a number of cells (N=100) simultaneously, with diffusion on the landscape dispersing the cells. These cells will then assume different fate outcomes, and we can calculate the penetrance by calculating the percentage of cells assuming a particular fate. In summary, our model outputs penetrance on E-to-MS fate differentiation when the POP-1 and SYS-1 levels are given.

### Connecting genotypes to phenotypes

Now we can calculate fate penetrance given POP-1 and SYS-1 levels, we need to connect mutant genotypes to these levels. The POP-1 and SYS-1 levels in each mutant will eventually come from the penetrance data by a model training process, but we can facilitate this process by incorporating the knowledge of the gene network and genetics. We incorporate the knowledge from gene network by imposing the directions of changes in POP-1 and SYS-1 levels, and to close the model we infer the POP-1 and SYS-1 levels in double mutants from the constituent single mutants.

First, we apply qualitative constraints on which direction a gene affects POP-1 and SYS-1 levels by our knowledge from the gene network. This can be achieved by following the gene network after the loss of function on the specific gene. For example, a mutation in *mom-4* will decrease WRM-1 and LIT-1, which in turn increases POP-1, while it has no known effect on SYS-1 level. Then we can only consider upward changes in POP-1 and constrain the change of SYS-1 to be zero. We can infer the directions of changes for other genes in a similar fashion, and a summary of these constraints is recorded in Table S1. The directions of changes in POP-1 and SYS-1 levels are then constrained when the model fits to the training dataset, making the training process more efficient.

Second, we can infer the POP-1 and SYS-1 levels for double mutants from the constituent single mutants. Not only would this decrease the number of parameters in the model, but it would allow the model to predict the penetrance of novel double mutants from combining existing single mutants in the model. For this purpose, a rule to determine double mutant penetrance from single mutant penetrance is required. While the most meticulous way is to write out the Michaelis-Menten equation for each reaction, this approach would make the model rely heavily on the known gene network, introduce additional parameters, and the model may lose the power to predict unknown interactions. Instead, we take a multiplicative model which is exact for SYS-1 and approximate for POP-1 under high POP-1 concentration in wildtype EMS cell. The fold-change of the POP-1 level relative to WT of a double mutant would be the product of factors for the two single mutations, and similarly for SYS-1 level (see Methods, Eqns. (3-4)). Therefore, the POP-1 and SYS-1 levels of double mutants are determined by those of single mutants, and predictions on penetrance can be made for double mutants with single mutations from the current data set, with no additional parameters.

### Parameters, fitting and predicting

For the model to describe the existing fate penetrance data and make falsifiable predictions, we need to train and validate the model by the data. First we introduce the parameters in the model. There are 2 types of parameters: general and gene-specific. The general parameters are those that are independent of the genotype, such as diffusion on cell fate and wildtype POP-1, SYS-1 levels, and the gene-specific parameters are those that only affect the landscape of embryos with a given genotype, through their effect on the POP-1, SYS-1 levels. A full list of parameters is recorded in Table S2 and their roles are described in the Methods.

To train the model by data, we need to find an optimal set of model parameters that fit the input data of mutants and their penetrance. The training process can be described in 2 steps: a grid search for gene-specific parameters and wildtype POP-1, SYS-1 levels, and a heuristic search for other general parameters. When fitting a set of double mutants, that all have one factor in common, such as the case of *sys-1(RNAi)* below, we find all parameters for the common factor that will work with all of the unique factors, and use the parameter range to assign error bars for the common factor. Since the training data is small, it is expected that there will be many solutions, and the challenge of fitting the model lies in charting the distribution of solutions. For this purpose, it is feasible to use a grid search method for gene-specific parameters since they are not involved in too many experimental constraints (see Methods).

Now we need to fit the general parameters. The grid search method can be extended to wildtype POP-1, SYS-1 levels, since there is an analytic formula for the dependence of fate on POP-1, SYS-1 levels. As many sets of general parameters can constitute solutions with proper gene-specific parameters, we used a heuristic search to find the general parameters that have the most tolerance on wildtype POP-1, SYS-1 levels, i.e. maximal area of *P*, *S* on the fate heat map. Perturbations in all 6 general parameters other than wildtype levels (see Methods) show that our choice of these parameters indeed has the most tolerance at least locally (Table S3).

To predict new double mutant penetrance from combinations of single mutations in the training dataset, the model simply draws all the permissible changes in POP-1, SYS-1 levels in these single mutants from the training process, and calculates the double mutant penetrance from all combinations of these changes. This process allows the model to not only predict the double mutant penetrance, but also provided an error bar on this prediction. In summary, after the model has been trained, it can predict double mutant penetrance and give an estimate of prediction error.

### Testing and validation

To draw quantitative conclusions from the model, we need to first fit and validate the model by the experimental data (Table 1). We partition the available data into a training and a testing set. As the training dataset cannot be randomly chosen, since an arbitrary subset of the data would not define all parameters, we describe how we choose the training dataset. First, we include the penetrance of all single mutants, since there is no way of characterizing them other than with data. Then we include all the double mutants with *sys-1(RNAi)* to constrain the *sys-1(RNAi)* parameters, since the penetrance for that mutant alone is close to zero. The data on synergistic interaction between *lit-1(t1534)* and *sys-1(RNAi)* also constrains the shape of the heat map controlled by general parameters. For similar reasons we include the double mutants between *mom-4* and *mom-5* to constrain the model, and we use the rest of the data as the testing dataset. Out of 36 data points on penetrance, 31 (including wildtype) are used for training, and 5 used for testing (Table 1).

After training, the model successfully fits the E-to-MS fate penetrance of the training dataset (Fig. 3A, Table S5), quantifies the changes in POP-1, SYS-1 levels by single mutations along with reasonable general landscape parameters (Fig. 3B, Table 3). Most data in the literature address the fate change in the E cell toward MS fate (Table 1), so in this work we focus on the E-to-MS fate differentiation and ignore other differentiations, such as MS-to-E and EMS self-renewal. The error bars come from both the variability of gene-specific parameters when the general parameters are fixed, and the variability of the general parameters. Thus the model successfully fits the training dataset and quantifies the gene-specific parameters.

**Fig. 3:**
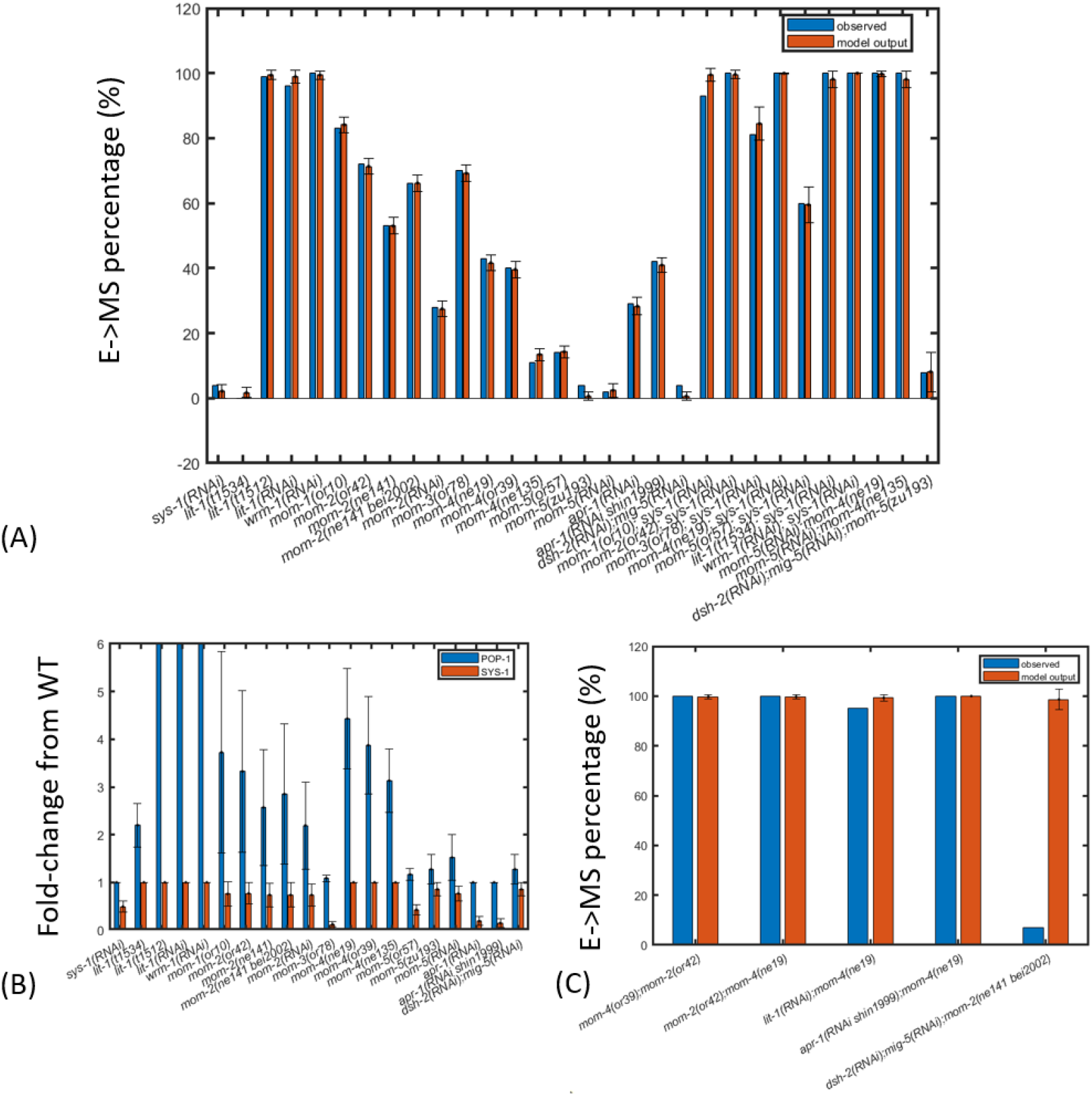
The landscape model is trained and validated by data on fate penetrance (see Table 1 for data and references). (A) The model successfully fit to all the training data with an error tolerance of for 5% and 10% error for single and double mutants respectively. In all panels in this figure, the error bar includes variations from both gene-specific and general parameters. (B) The model is trained by fitting the fold-changes relative to WT caused by the single mutations, and the model explores the admissible fold-changes constrained by the training dataset. Large variations tend to occur in POP-1 changes as the model does not set an upper bound on the POP-1 level, which biologically comes from the POP-1 level in EMS (see Methods). (C) The model correctly predicts all but one testing data point. Most double mutants in the testing data exhibit full penetrance (Table 1), so only provide constraints on parameters rather than equalities as would be the case with partial penetrance. The only discrepancy is biologically perplexing as already noted in the original reference (Bei et al., 2002).

The landscape model correctly predicts the testing dataset except for one mutant penetrance (Fig. 3C). The only discrepancy comes from the *dsh-2(RNAi); mig-5(RNAi); mom-2(ne141)* triple mutant, where the model predicts a much higher penetrance than observed (Fig. 3C). It is not surprising that the model overestimates the penetrance, as *dsh-2(RNAi)*, *mig-5(RNAi)*, and *mom-2(ne141)* change POP-1, SYS-1 levels in the same direction from the gene network (Fig. 1C, Table S1), so the penetrance of *dsh-2(RNAi); mig-5(RNAi); mom-2(ne141)* should at least be higher than that of *mom-2(ne141)*. This is clearly not the case from the data (Table 1), and this effect was attributed to the “independent function of Disheveled” in the original work (Bei et al., 2002). Since this observation is not consistent with the known gene network, any model based on the known gene network cannot possibly predict this observation. In summary, the model successfully fits to the training dataset, and predicts all testing dataset except for one datapoint that does not agree with the gene network.

### The model quantifies POP-1 and SYS-1 levels in wildtype and mutant embryos

The model quantifies the developmental landscape in several respects: (1) the model reveals the nonlinear response of fate penetrance on POP-1 and SYS-1 levels. (2) The model quantifies the tolerance of wildtype embryos to variations in POP-1 and SYS-1 levels. (3) The model quantifies the changes in POP-1 and SYS-1 levels in single mutants.

First, the model reveals the nonlinear response by generating a fate heat map directly from Wnt Michaelis-Menten dynamics. Given a set of general parameters, the model generates a fate heat map by calculating the fate penetrance on a grid of POP-1 and SYS-1 levels (Fig. 4A). This fate heat map demonstrates an unexpected dependence of fate response on POP-1 and SYS-1 levels: the fate contours are concave, which implies that cell fate is not simply a function of the POP-1/SYS-1 ratio.

**Fig. 4:**
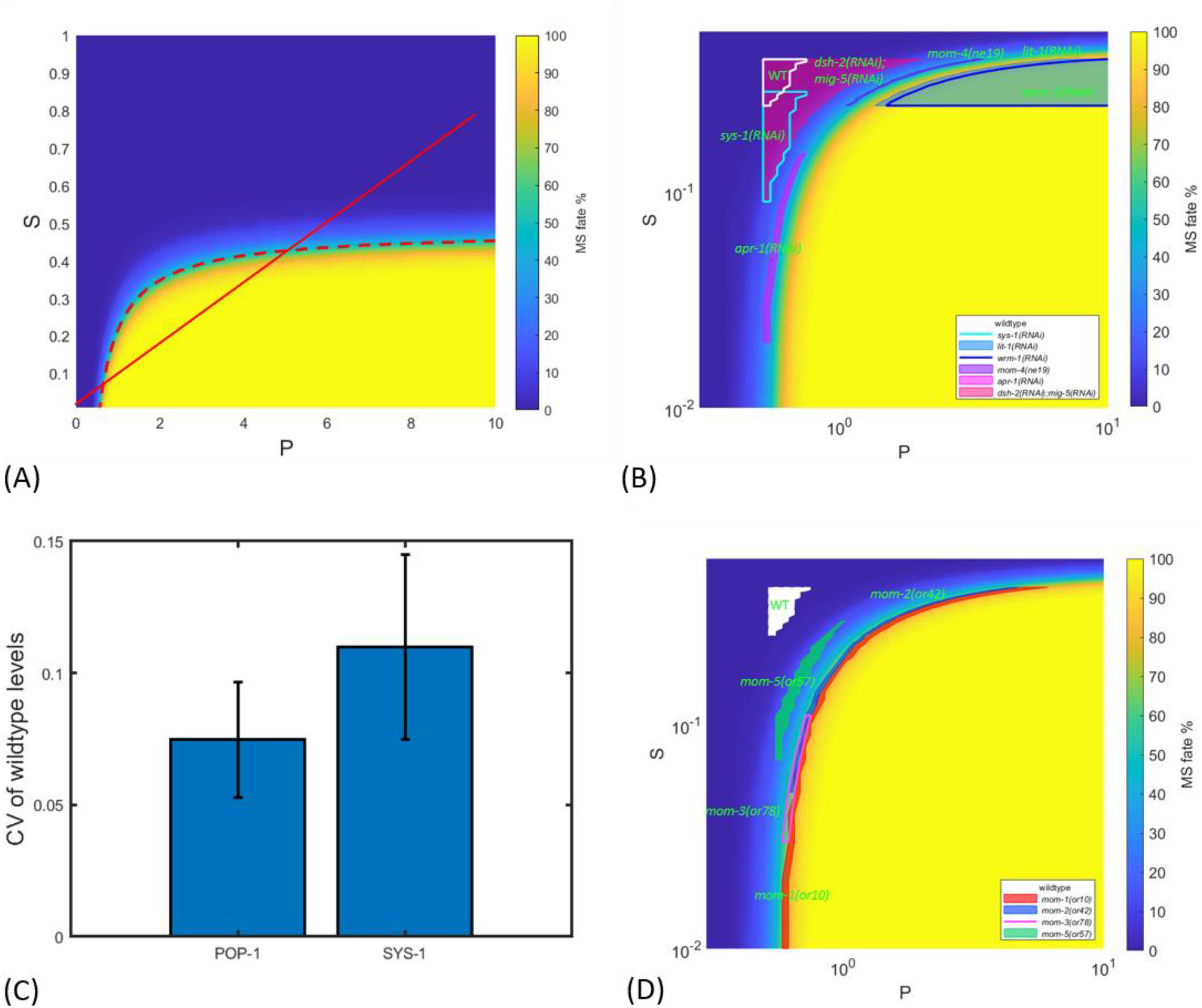
The model quantifies the landscape in EMS development. (A) The shape of the fate heat map is quantified by the Wnt Michaelis-Menten dynamics. The model generates the fate heat map by calculating the penetrance of E->MS fate differentiation for each POP-1 (P) and SYS-1 (S) level. The equi-penetrance contours of the fate heat map are concave (red dashed curve: contour for 50% penetrance). Contrary to prior understanding (Huang et al., 2007), cell fate changes dramatically for fixed POP-1/SYS-1 ratio (red solid line). (B) The model maps out the regions of fate heat map for wildtype embryos and single mutants. The regions are determined by both the training dataset of fate penetrance and the constraints from gene network (see Methods). The regions reveal correlation between POP-1 and SYS-1 levels imposed by the training dataset, which results in large error bars in individual POP-1 and SYS-1 levels in Fig. 3B, but tight constraints on penetrance. The panel is generated under fixed generic parameters except for wildtype POP-1 and SYS-1 levels. Clarification: *wrm-1(RNAi)* largely overlaps with *lit-1(RNAi)*, with the latter containing the former. We plot only selected single mutants for visibility (see Table S2 and Fig. S1 for all single mutants). (C) The model measures the wildtype tolerance on variability in POP-1 and SYS-1 under the training dataset. The variability is measured by Coefficient of Variation (CV), which is standard deviation divided by the mean, and the error bar is calculated from perturbing the 6 general parameters (see Methods). (D) The model puts Wnt upstream genes in different functional groups. *mom-1(or10)* has its region span across most of the heatmap. *mom-2(or42)* is constrained to the upper-right, and *mom-3(or78)* and *mom-5(or57)* are constrained to the lower-left of the heat map.

First, the concavity of the fate contours is apparent in the fate heat map and derived from Michaelis-Menten dynamics. For a fixed concentration ratio of POP-1 and SYS-1, the cell assumes E fate when both POP-1 and SYS-1 are close to zero, MS fate for small POP-1 and SYS-1, and E fate again for large POP-1 and SYS-1 (Fig. 4A). A bias toward E fate at low POP-1 and SYS-1 levels was determined by the totality of the fit to the data (see Methods and Table S2), but the change from MS fate to E fate when increasing POP-1 and SYS-1 while fixing the ratio was a direct consequence of just Wnt Michaelis-Menten dynamics. Under the assumption that POP-1 and SYS-1 levels are higher than the promoter levels, one can express the shape of contours in the plane of P,S (the POP-1 and SYS-1 levels) as follows:

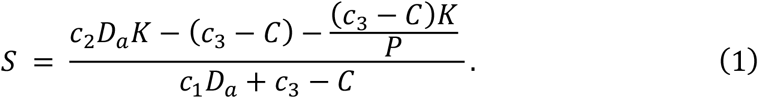

Here *C* is slope of the potential in Fig. 2 at x=0 which is a monotonic function of the penetrance and is related to P, S by solving the Michaelis-Menten equations. Nonzero penetrance requires *c*_3_ > *C*, the other parameters are positive, and P is constrained to keep S>0. (see Methods and Eqn. (5) for parameter definition and derivation of contour formula). The concavity of these contours indicates that an increase in POP-1 and SYS-1 levels with a fixed ratio will start to decrease the E->MS penetrance at certain POP-1, SYS-1 levels.

Second, the model quantifies the tolerance of wildtype embryos on POP-1 and SYS-1 variability. For each of 25 sets of general parameters, the model gives a range of wildtype POP-1, SYS-1 levels allowed by the training dataset (Fig. 4B; see Fig. S1 for all single mutants). We can quantify the variability by calculating the coefficient of variation (CV) in POP-1 and SYS-1 levels over those 25 sets of parameters. Both CVs in POP-1 and SYS-1 are around 1/10 (Fig. 4C), which means their wildtype levels are constrained by the penetrance data. This result shows that the wildtype POP-1 and SYS-1 levels can be reliably inferred from the penetrance data.

If the only data we had for a given mutant allele was its penetrance, then all we could say is that P,S would be restricted to the appropriate contour in the fate plane. The allowed region would be broadened further by whatever uncertainty the fit allows for the general parameters. Further constraints come the genetic network Fig 1C. For instance, mutations in genes that activate SYS-1 can only lower its value below WT not augment it. The same hold for RNAi. The underlying concavity of the fate heat map imposes strong correlations on the errors in P and S, Fig. 4B. Thus domain of *sys-1(RNAi)* is broad in S but quite constrained in P.

Further restrictions on the P,S domains of mutants in Fig. 4B come from double mutants but their consequences can be subtle, again due to the underlying concavity of the fate heat map. To illustrate, consider the *mom-2(or42)* and *mom-3(or78)* alleles that both have a penetrance around 70%. When we combine each of these with *sys-1(RNAi)* the former is fully penetrant and the latter only 81% penetrant. Since we know SYS-1 can only be lowered in the double mutant, the fit places the *mom-2* allele largely in the region P > WT where the contour is more horizontal, so moving it modestly downward will place it somewhere in the yellow region. (Precisely where we cannot tell since the penetrance is full). The penetrance of *mom-3(or78)* increases only slightly when combined with *sys-1(RNAi)*, thus it is placed more in the vertical part of the contour, with P ~ WT and S much smaller than WT. So the double mutant will lower SYS-1 further than in *mom-3(or78)* alone, and change the penetrance only slightly. Our training data in fact includes in total, 7 mutants *wrm-1(RNAi), mom-1(or10), mom-2(or42), mom-3(or78), mom-4(ne19), mom-5(or57),* and *lit-1(t1534)* (ordered by decreasing penetrance) all of which were subject to *sys-1(RNAi)* in a consistent way. Four of them are fully penetrant with *sys-1(RNAi)*, so tend to be placed in the region P > WT. *mom-5(or57)* is interesting since its penetrance changes substantially from 14% in the single mutant to 60% in the double. The fit gives it comparable variation along the two axes.

Thus, the most informative data for the fit, are multiple alleles with low penetrance that becomes larger but still less than full when combined with common second mutation. Or conversely a large penetrance set that all become lower in a double mutant. The variability of RNAi works to our advantage provided its applied consistently to each allele. Each protocol functions like a new allele. Fluorescently tagged POP-1 and SYS-1 would be the most immediate way to test our predictions. But note, much sharper predictions are made when both species are measured because of the dominant constraint of the concave contours in the fate density map.

### The model predicts fate response in antagonistic genetic interactions

Our model has the ability to quantitatively predict fate differentiation of double mutants from single mutants in the data. The double mutant penetrance can be predicted in the same way the testing data are predicted (see Methods). The gene network allows us to qualitatively infer the fate differentiation, but our model makes the quantitative prediction possible. This is especially helpful when the genes involved act in opposite directions on a downstream target, as there can be a wide range of potential outcomes. In this section we show unintuitive model predictions on mutants in *pop-1* branch genes that can be experimentally tested.

All the single mutants in our data act in the same direction to increase POP-1 and decrease SYS-1 levels (Table S1). Therefore, pop1 mutants (or *sys-1* gain-of-function mutants) are antagonistic to what we have and therefore useful test of model.

The *pop-1* mutant alone would tend to induce MS→E but should not cause E→MS. However in Fig. 5A and Table S4, we predict the change in penetrance for multiple alleles that our fit places in the region P > WT. A 25% change is most informative since a 50% reduction gives nearly zero conversion of E to MS. We define the fold change to be the penetrance for the single allele divided by the penetrance when combined with a 25% reduction in POP-1. As expected the fold change is generally less for alleles with higher POP-1 levels. This implies that in general mutants in the *pop-1* branch of Fig. 1C will be less affected by a reduction in pop-1 than those in Wnt upstream or *sys-1* branches. The most extreme instance is *apr-1(RNAi)* whose fold change is 25, since it occupies the lower left corner of the density plot where the contour lines are nearly vertical. However the fold change can differ by a factor of two for different alleles of the same gene, e.g., *mom-2*, largely due to difference in single mutant penetrance which leads to these alleles occupying different regions on the heat map.

**Fig. 5:**
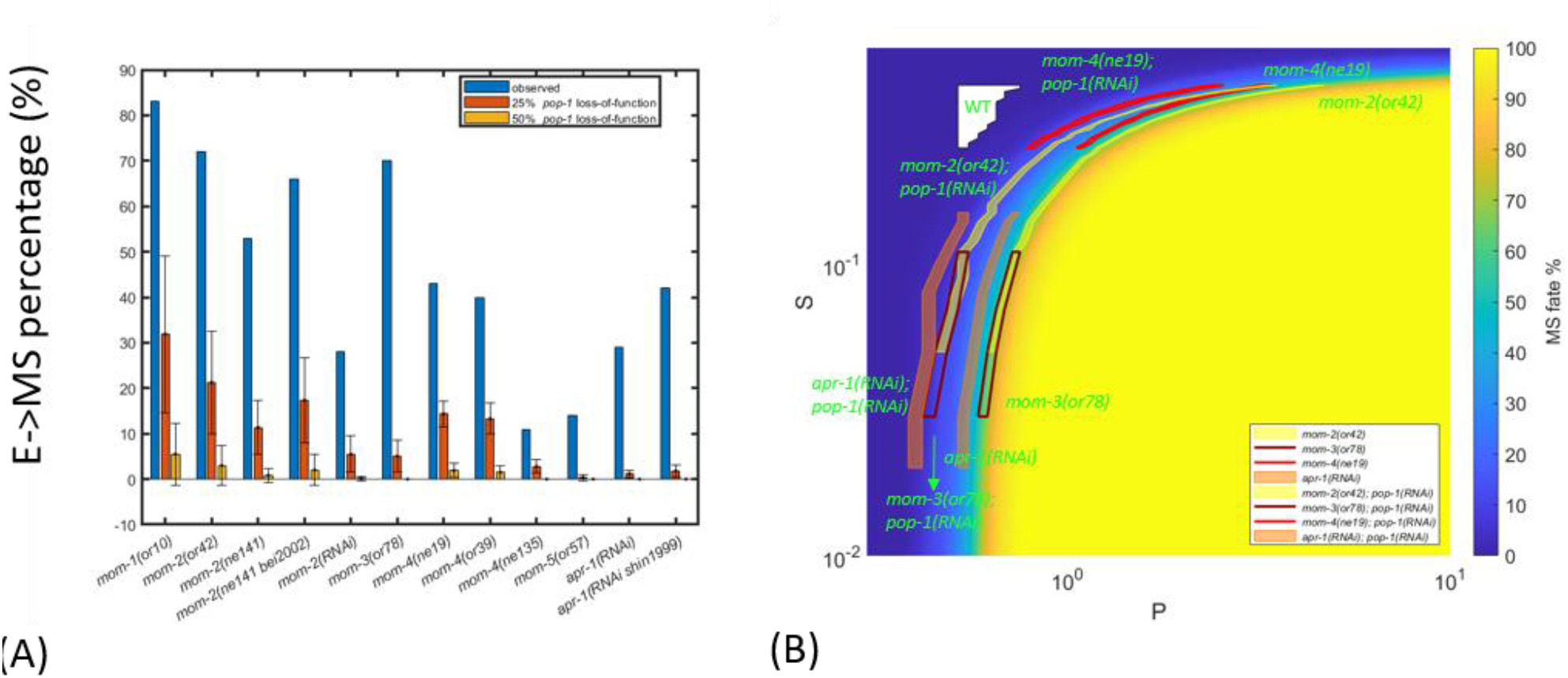
The model predicts fate response for antagonistic genetic interactions. Antagonistic genetic interactions indicate that one mutation works in the opposite direction to another mutation on downstream genes. Here the *pop-1* loss-of-function acts against other single mutants in POP-1 level. (A) The model predicts the penetrance of single mutants in the data under the *pop-1* loss-of-function background, such as RNAi interference. Shown: single mutant penetrance under 0%, 25%, and 50% reduction in POP-1 level. (B) The change of POP-1 and SYS-1 levels under the *pop-1* loss-of-function background with a 25% reduction in POP-1 level explains the model predictions. Mutants with and without *pop-1* loss-of-function have the same color.

The second unexpected prediction of the model comes from the non-monotonicity among Wnt upstream genes. The model places Wnt upstream genes in different functional groups based on the model’s interpretations of double mutant penetrance data, which manifests itself when the model predicts double mutant penetrance. The strongest result again comes from comparing *mom-2(or42)* and *mom-3(or78)*, which have similar penetrance in single mutants. Under a 25% reduction in POP-1 level, the *mom-2(or42)* has a fold-change of 3.3 after *pop-1* loss-of-function, higher than 14 of *mom-3(or78)* (*p* < 10^−4^, one-sided bootstrapping). The reasoning follows the same argument above as *mom-3(or78)* has a lower POP-1 level. Notably the penetrance of *mom-2(or42)* under POP-1 reduction has a large error bar, which comes from the wide POP-1 level enclosed in the region of *mom-2(or42)* single mutant (Fig. 5B). This can be narrowed down by further double mutant experiments. In summary, the model predicts varied penetrance response of Wnt upstream genes under the *pop-1* loss-of-function background.

## Conclusions and Discussions

In this work we built a quantitative model for cell fate differentiation by including Wnt dynamics into Waddington’s developmental landscape. Our model is successfully trained and validated by existing data on fate penetrance, and quantifies the developmental landscape in several aspects. The model reveals the nonlinear fate dependence on POP-1/TCF and SYS-1/beta-catenin. The model also translates fate penetrance into POP-1 and SYS-1 levels, and explains counter-intuitive double mutant fate penetrance results. Finally, the model gives falsifiable predictions on antagonistic gene interactions, which would be difficult to make by gene network models.

Waddington’s landscape is a suitable model for describing cell fate decisions, as the model typically contains few parameters, and they are closely related to the developmental process. There are 8 general parameters and 2 gene-specific parameters (POP-1 and SYS-1) for each genotype, compared to thousands of parameters required to build an artificial neural network for modeling dynamical processes (Yazdani et al., 2020). These parameters also have intuitive interpretations, which allow us to predict biologically meaningful quantities such as POP-1 and SYS-1 levels in mutants. We expect the landscape model to play an important role in modeling developmental processes.

The model reveals nonlinear fate dependence on the POP-1/SYS-1 ratio, which can be experimentally verified. It is clear from the gene network for Wnt pathway that qualitatively speaking high POP-1 and low SYS-1 levels lead to MS fate, while low POP-1 and high SYS-1 levels lead to E fate. However, how fate response quantitatively changes with POP-1 and SYS-1 has not been clear. Our results show that, under the same POP-1/SYS-1 ratio, the E cell would admit MS fate at low POP-1 level and E fate at high POP-1 level. To verify this prediction, one can imagine combining alleles of genes in pop-1 or sys-1 branches with different strengths, and measuring POP-1 and SYS-1 levels by quantitative measurements from live imaging (Zacharias et al., 2015). These experiments can deepen our understanding of the Wnt dynamics.

Our model can be generalized to quantify other fate differentiations in EMS development, such as E-to-C fate change and EMS self-renewal. Our methodology can already be applied in its present form to the MS cell, but there is insufficient data to fit. Finally, our model can describe EMS self-renewal, which is a recently discovered phenotype that describes MS-to-EMS fate differentiation (Du et al., 2014; Du et al., 2015). Fate reiteration is already incorporated in the landscape model, as parameters related to SKN-1 decay control the quadratic-to-quartic transition in the landscape potential, and delaying this transition keeps the cells in a region of EMS fate. It will be interesting to see if the model can incorporate double mutants with EMS self-renewal when the data are available. To add the C fate requires a three way decision which would likely necessitate a potential in two dimensions with sectors in the plane allocated to the three fates. Genes related to the C fate, such as *pal-1*, will also need to be considered (Hunter and Kenyon, 1996).

## Author Contribution

SSC, ZB, and EDS wrote and reviewed the paper. SSC did the model calculation.

## Acknowledgements

This study was partly supported by an NIH grant (R01GM097576) to Z.B. and an NSF grant PHY 2013131 to EDS. Research in Z.B. lab is also supported by an NIH center grant to MSKCC (P30CA008748).

## Methods

### Model

#### Wnt Michaelis-Menten Dynamics

To connect POP-1 and SYS-1 levels to cell fate, we model the binding of these transcription factors to promoters of fate genes by the Michaelis-Menten Dynamics (Fig. 2B). The full equations are as follows:

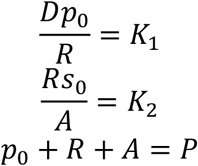

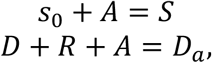

where *D*, *R*, *A* are the concentration of unbounded, repressive, and activate forms of promoter, *p*_0_, *s*_0_ are the concentration of unbounded POP-1, SYS-1, *P, S* are the total concentration of POP-1, SYS-1 in the cell nucleus, *K*_1_, *K*_2_ are the equilibrium constants of the chemical reactions in Fig. 2B, and *D*_*a*_ is the total concentration of promoters. After non-dimensionalizing the unit of concentration by *K*_1_ we have

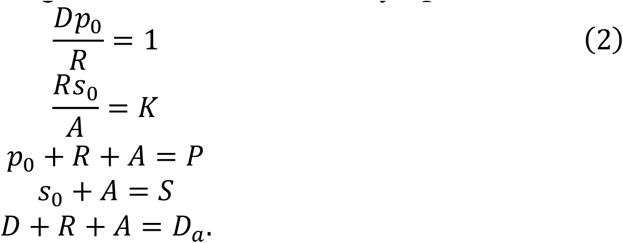

These equations can be solved for *p*_0_, *s*_0_, *D*, *R*, *A* if *P*, *S*, *K*, *D*_*a*_ are given. As each single mutation has its own *P, S* parameters to be fit to the training data set, the Wnt Michaelis-Menten dynamics has 2 undetermined parameters, *K* and *D*_*a*_. The ratio of equilibrium constants *K* determines the relative binding strength of POP-1 and SYS-1 to the promoter complex, and changing this constant would affect e.g. the amount of SYS-1 relative to POP-1 in order to activate E fate. Hence *K* can only be decided by observing fate outcomes while changing POP-1 and SYS-1 in a quantitative fashion, and here we simply assume *K* = 1, meaning no bias on the relative binding strength. *D*_*a*_ represents the amount of promoter, and we assume that *D*_*a*_ < *P, S* in our model. These equations can then be solved by symbolic equation solvers, and only one of the 4 solutions is physical when imposing the parameter range *S* ∈ [0,1], *P* ∈ [0,10].

#### Inferring POP-1 and SYS-1 levels of double mutants from single mutants

To predict double mutant penetrance from two single mutants A, B with fitted POP-1, SYS-1 levels, the model requires a rule to infer POP-1, SYS-1 levels of double mutants from those of its single mutations. It is not practical to attempt to model the entire gene network relating A,B so instead we resort to a product rule:

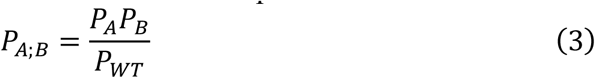

where *P*_*A;B*_, *P*_*A*_, *P*_*B*_, *P*_*WT*_ are the POP-1 levels of double mutant *A*; *B*;, *A*, *B*, and wildtype in E cell. This is plausible when the levels of the mutants are well below WT levels so that the species being compared are limiting. A clear problem with this assumption arises during EMS cell division, POP-1 in EMS decreases in the future E cell nucleus (Sawa and Korswagen, 2013), which imposes a hard bound on POP-1 in the E cell.

On the other hand, SYS-1 levels in EMS and E cells are comparable, and gene perturbations decrease SYS-1 in E cell (Sawa and Korswagen, 2013). Thus the corresponding rule for SYS-1 levels in double mutants is more plausible:

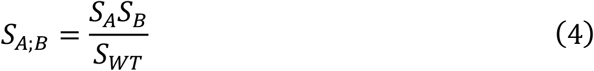

where *S*_*A;B*_, *S*_*A*_, *S*_*B*_, *S*_*WT*_ are the SYS-1 levels of double mutant *A*; *B*, *A*, *B*, and wildtype in E cell.

#### Deriving penetrance from landscape model

The quantitative landscape model determines cell fate penetrance by moving cells according to the landscape potential. This dynamic is solved for t∈ [0,2], with *t* ∈ [0,1) and *t* ∈ [1,2] representing EMS and MS, E cell life respectively. The potential takes the form:

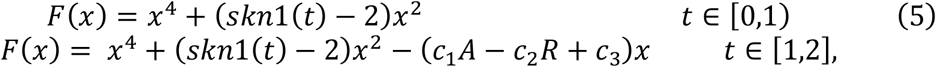

where

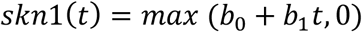

represents the decay of SKN-1, which is modeled by a linear decay with *b*_0_ > 0, *b*_1_ < 0. The decrease in quadratic coefficient turns the single-well potential into a double-well potential (Fig. 2A), which unlocks the MS and E fates. Right after EMS division (*t* = 1), the Wnt signal biases the landscape potential by changing the linear term, with repressive and activate forms of promoter solved from the Wnt Michaelis-Menten dynamics. We impose the constraints *c*_1_, *c*_2_ > 0 (and *A, R* > 0) so that active promoters (*A*) lead to E fate, while repressive promoters (*R*) lead to MS fate. *c*_3_ can be of either sign, but as *c*_3_ > 0 corresponds to E fate in wildtype E cells, it is our starting point when searching for general parameters. *N* = 100 cells then follow the potential in *t* ∈ [0,2], and cell fates are determined by their terminal positions when *t* = 2. Specifically a cell assumes MS fate when *x*(2) < −0.2, E fate when *x*(2) > 0.2, and EMS fate when −0.2 ≤ *x*(2) ≤ 0.2, and the penetrance of a certain fate is the percentage of cells assuming the fate. The boundary 0.2 was manually selected, and it does not affect the fit. When −0.2 ≤ *x*(2) ≤ 0.2, the model predicts EMS self-renewal, which has been reported in previous work (Du et al., 2014; Du et al., 2015). The cell fate trajectory is simulated by the discretized stochastic differential equation:

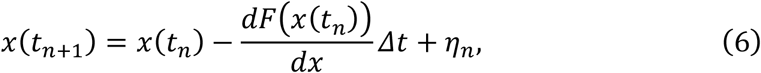

where *Δt* = 0.02 and *η*_*n*_ is sampled from a Gaussian distribution with mean 0 and variance 4*DΔt*, where *D* is the diffusion constant on the fate coordinate. The landscape dynamic can then be simulated given 8 general parameters: *b*_0_ − *b*_1_, the intercept and slope for SKN-1 decay, *D*, the diffusion of cell fate on the landscape, *c*_1_ − *c*_3_, the parameters relating levels of activated and repressed promoters (*A, R*) to the landscape, and *P*, *S*, the POP-1 and SYS-1 levels in wildtype E cells, which are fit to the training data set (see Methods)

### Model training and testing

#### Training process

To train the model to fit the training dataset, we split the training process into 2 steps: (1) a grid search method for finding all admissible gene-specific parameters and wildtype POP-1, SYS-1 levels with assumed general parameters, and (2) a heuristic method for finding the remaining 6 general parameters. The grid search being exhaustive allows for correlations among the fitting parameters.

The grid search method is carried out over 3 steps: (1) calculating the fate heat map under a fixed set of general parameters other than wildtype POP-1, SYS-1 levels, (2) finding fate contour for each single mutant from the single mutant penetrance, and (3) trimming these fate contours by the double mutant penetrance.

To begin with, we calculate the fate heat map for a range of POP-1 and SYS-1 values. We use a range of [0,1] for S and [0,10] for P, with a linear resolution of 0.01, 0.033 respectively. Then we calculate the fate penetrance for each of the grid points. This fate heat map is then used for the following steps.

Then we find the fate contour for each single mutant by their own penetrance. Given a single mutant penetrance, we can find all grid points in P, S with fate penetrance close enough to the observed penetrance within a tolerance level (we used 5%). In this step the training of each single mutant only depends on their own observed penetrance.

Finally we use the information from double mutants to trim down the fate contours for each single mutant involved. For each observed double mutant penetrance, we look at all the combinations of allowed grid points from single mutant contours, calculate the double mutant penetrance from the rule we introduced earlier in the Methods, and keep or discard this combination of grid point values based on whether the calculated double mutant penetrance is close enough to the observed one within a tolerance level (we used 10%). The situation is more complicated when one single mutant is involved in several observed double mutant penetrance, e.g. *sys-1(RNAi)*. In this case we loop through all grid points in the fate contour of the common single mutant, test the combinations with all other single mutants first, and then only include a grid point of the common single mutant when this grid point allows at least one solution in all the observed double mutant penetrance involved. After this process, for each set of general parameters we can get a list of admissible values for each single mutant.

One can see we can easily find all wildtype POP-1, SYS-1 levels that allow for at least one solution of all gene-specific parameters: all we need to do is to repeat the process for each grid point of wildtype POP-1, SYS-1 levels. This is computationally plausible as we do not need to recalculate the heat map for different wildtype POP-1, SYS-1 levels, which is the case for other 6 general parameters. Then we have a list of admissible wildtype POP-1, SYS-1 levels.

In principle we can do a full grid search for the other 6 general parameters; however a fate heat map needs to be calculated when we change any of these parameters, which greatly slows down the algorithm. Also it is not clear what criteria we should use to select for these parameters: most general parameter sets around the parameter set we use have more than 10 wildtype POP-1, SYS-1 level solutions (Table S3). For almost all 6 parameters other than wildtype POP-1, SYS-1 levels, the number of wildtype POP-1, SYS-1 solutions generally peaks at 0% perturbation and decreases with the strength of perturbation. The only exception is the SKN-1 intercept, where a small number of solutions occurs at 5% perturbation. This is due to the effect of SKN-1 intercept on the gradient of the heat map, especially the slope of POP-1 on the lower-right. For example, high precision is required for fitting the *mom-3(or78); sys-1(RNAi)* penetrance data, as the SYS-1 level of *mom-3(or78)* travels through a region of large slope in POP-1 level under the *sys-1(RNAi)* background. Since SKN-1 intercept affects this slope in POP-1 level, the density of the grid limits the number of solutions we found. We decided to score the general parameter sets by the number of wildtype POP-1, SYS-1 level solutions. First, we used a gradient descent approach, with the target function being the sum of squared difference in the calculated and observed penetrance, to find one set of general parameters that gives a solution. Then we did a grid search for the 6 parameters for those that give maximal area in POP-1 and SYS-1 levels which fit all training data. Eventually we found a set of general parameters that satisfies this criterium at least locally (Table S3).

#### Testing process

To validate our model against a testing dataset, we need to make predictions on the mutant penetrance in the testing dataset based on the parameters we found in the training process. For each set of general parameters, we have a list of admissible gene-specific parameters for each single mutant. When we want to predict a double mutant that is a combination of 2 single mutants in the training dataset, we can simply consider all possible combinations from the 2 lists of gene-specific parameters and calculate the penetrance. The average penetrance from all possible combinations is then used as the model prediction.

#### Error bar calculation

To know how reliable our model estimates biological parameters and predicts fate penetrance, we need to have a notion of error bar on our estimations. There are 2 sources of variations on our model: variations on gene-specific and general parameters. The variation on gene-specific parameters is built into the training process of the model, and our grid search method finds all admissible gene-specific parameters compatible with the training data. The variation on general parameters comes from uncertainties on estimations of these parameters, as there are many sets of general parameters that are compatible with the training data. To quantify this uncertainty, we perturb the 6 general parameters other than wildtype POP-1 and SYS-1 levels by a fixed amount (10%), and attempt to find solutions under the perturbed general parameters. All perturbations are compatible with the training dataset, i.e. they all admit at least one solution, but they contain different numbers of solutions in wildtype POP-1 and SYS-1 levels (Table S3). Then we include a 6% variation on general parameters by considering all the perturbations when estimating parameters and predicting fate penetrance.

### Model quantification of the developmental landscape

#### Derivation of contour formula on fate heat map of POP-1 and SYS-1 levels

To understand how the cell fate changes with POP-1 and SYS-1 levels, here we derive the formula for the contour levels in the fate heat map. We start with the simplified Michaelis-Menten equations:

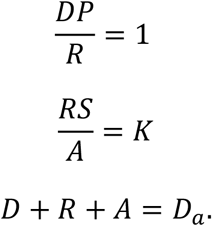

We use the *P, S* >> *D*_*a*_ approximation from the assumption that POP-1 and SYS-1 are more abundant than promoters. We can solve for activated and repressed promoter levels, *A, R*, from these equations:

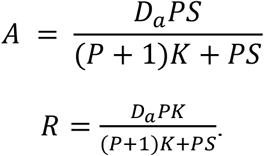

The fate contours can be expressed by Eqn. (5):

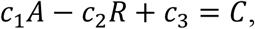

where *C* is the coefficient of x in landscape potential. *C* is tightly connected to fate penetrance, as the landscape potential determines penetrance up to dispersion in cell fate. *C* > 0 corresponds to zero penetrance, which is of no physical interests. Hence we can assume *C* < 0 and that *c*_3_ − *C* > 0 (*c*_3_ > 0 as the wildtype fate of E cell is E fate). Plugging in the expressions for *A, R* we obtain

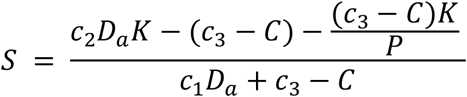

as Eqn. (1) in the main text. *c*_2_*D*_*a*_*K* is the maximal value that *c*_2_*R* can take when *K* = 1, and since *c*_2_*R* − (*c*_3_ − *C*) = *c*_1_*A* > 0, we have *c*_2_*D*_*a*_*K* − (*c*_3_ − *C*) > 0, and that for any contour with level *C* there is a POP-1 value *P*_*C*_ > 0 such that *S* = 0, which is the starting point of the contour on the *P*-axis. Then *S* increases with *P* concavely until *S* saturates at 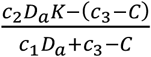.

