## Supplemental Figure 1 and Supplemental Table 1-5 for "A landscape model for cell fate decisions during mesoendoderm differentiation in C. elegans based on Wnt dynamics"

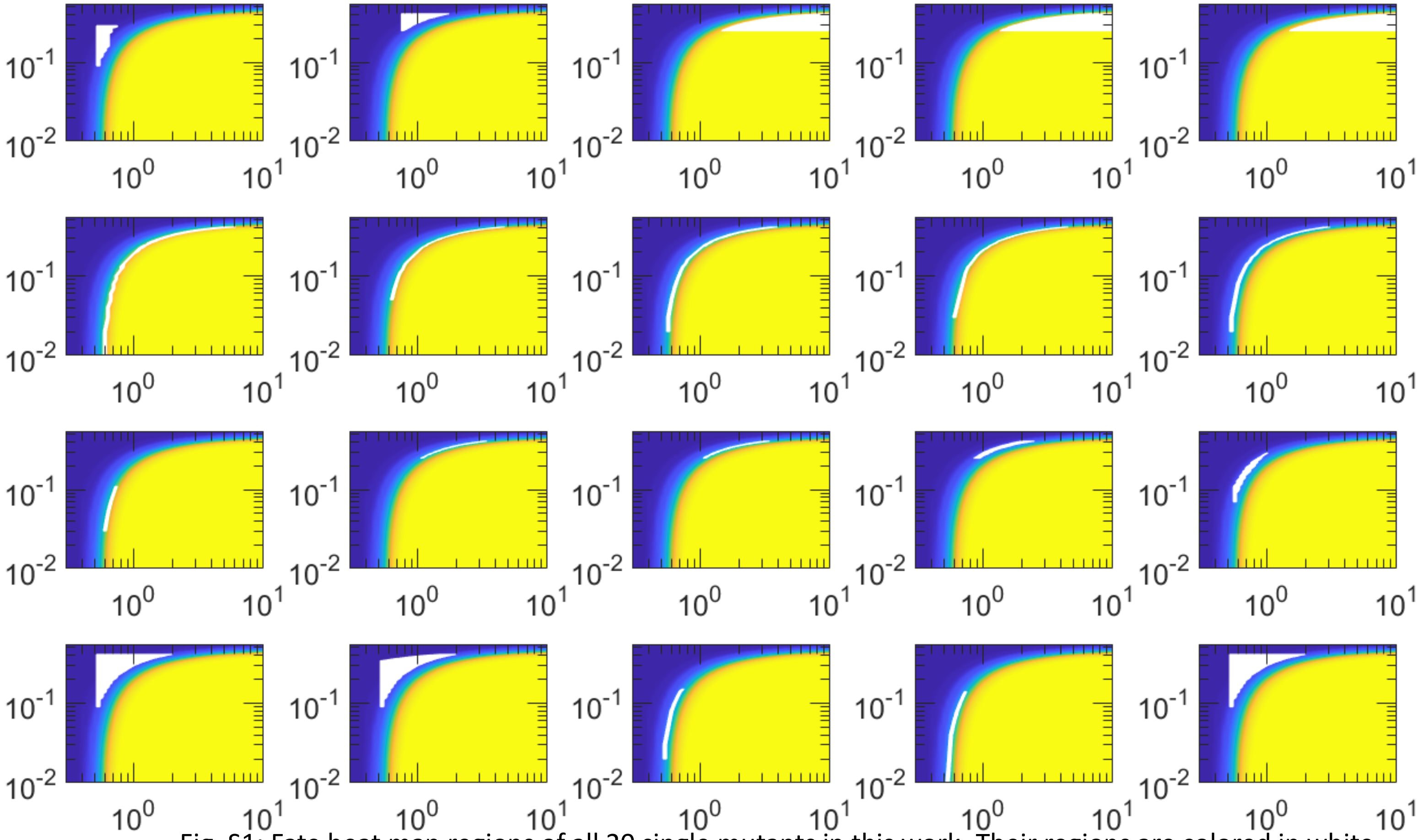


Fig. S1: Fate heat map regions of all 20 single mutants in this work. Their regions are colored in white, and the greet texts indicate the single mutants. The color map represents fate penetrance as in Fig. 4B. The single mutants are, from left to right followed by up to down:

*sys-1(RNAi), lit-1(t1534), lit-1(t1512), lit-1(RNAi), wrm-1(RNAi)*

*mom-1(or10), mom-2(or42), mom-2(ne141), mom-2(ne141 bei2002), mom-2(RNAi)*

*mom-3(or78), mom-4(ne19), mom-4(or39), mom-4(ne135), mom-5(or57)*

*mom-5(zu193) ,mom-5(RNAi), apr-1(RNAi), apr-1(RNAi shin1999), dsh-2(RNAi);mig-5(RNAi)*

| | Direction in change of levels | | | |
|---|---|---|---|---|
| | MS | | E | |
| | POP-1 | SYS-1 | POP-1 | SYS-1 |
| *sys-1(RNAi)* | unchanged | down | unchanged | down |
| *lit-1(t1534)* | up | unchanged | up | unchanged |
| *lit-1(t1512)* | up | unchanged | up | unchanged |
| *lit-1(RNAi)* | up | unchanged | up | unchanged |
| *wrm-1(RNAi)* | up | unchanged | up | unchanged |
| *mom-1(or10)* | down | down | up | down |
| *mom-2(or42)* | down | down | up | down |
| *mom-2(ne141)* | down | down | up | down |
| *mom-2(ne141 bei2002)* | down | down | up | down |
| *mom-2(RNAi)* | down | down | up | down |
| *mom-3(or78)* | down | down | up | down |
| *mom-4(ne19)* | up | unchanged | up | unchanged |
| *mom-4(or39)* | up | unchanged | up | unchanged |
| *mom-4(ne135)* | up | unchanged | up | unchanged |
| *mom-5(or57)* | down | down | up | down |
| *mom-5(zu193)* | down | down | up | down |
| *mom-5(RNAi)* | down | down | up | down |
| *apr-1(RNAi)* | down | unchanged | unchanged | down |
| *apr-1(RNAi shin1999)* | down | unchanged | unchanged | down |
| *dsh-2(RNAi);mig-5(RNAi)* | down | down | up | down |

Table S1: The constraints on changes in POP-1 and SYS-1 levels applied on each mutation in this work. Some constraints are less obvious from the gene network. For example, when Wnt upstream genes, *mom-1,2,3,5, dsh-2,* and *mig-5* are mutated, the SYS-1 level is affected in two ways: First, SYS-1 is no longer prevented from the degradation (Fig. 1C, arrow pointing from *dsh-2*, *mig-5* to *sys-1*) and its level decreases (Sawa and Korswagen, 2013). Second: There are more APR-1 in the posterior side of EMS (Fig. 1D), which increases SYS-1 level in E cell as APR-1 facilitates the prevention of SYS-1 degradation from Wnt upstream genes (Huang et al., 2007). However, experimental evidence points out that SYS-1 level in E cell increases in Wnt upstream mutants (Huang et al., 2007), so we put constraints on these genes accordingly. Finally, mutation in *apr-1* should also lead to decrease in POP-1 according to the gene network (Fig. 1C), but the effect should be small as most APR-1 concentrates in the anterior side of EMS, so we constrain the POP-1 level in E cell to be unchanged for *apr-1* mutants.

| general parameter | mean | SD (fixed) | | |
|---|---|---|---|---|
| SKN-1 slope | -6.30 | -0.50 | | |
| SKN-1 intercept | 7.00 | 0.55 | | |
| cell fate diffusion | 0.001 | 7.91E-05 | | |
| c_1 | 1.6E+04 | 1.26E+03 | | |
| c_2 | 1.6E+04 | 1.26E+03 | | |
| c_3 | 2.73 | 0.22 | | |
| genotype | P mean | P SD | S mean | S SD |
| wildtype | 0.59 | 0.06 | 0.35 | 0.05 |
| | P ratio mean | P ratio SD | S ratio mean | S ratio SD |
| *sys-1(RNAi)* | 1.00 | 0.00 | 0.49 | 0.12 |
| *lit-1(t1534)* | 2.20 | 0.45 | 1.00 | 0.00 |
| *lit-1(t1512)* | 11.32 | 3.84 | 1.00 | 0.00 |
| *lit-1(RNAi)* | 11.08 | 3.87 | 1.00 | 0.00 |
| *wrm-1(RNAi)* | 12.12 | 3.79 | 1.00 | 0.00 |
| *mom-1(or10)* | 3.73 | 2.11 | 0.76 | 0.25 |
| *mom-2(or42)* | 3.33 | 1.69 | 0.77 | 0.22 |
| *mom-2(ne141)* | 2.57 | 1.22 | 0.74 | 0.25 |
| *mom-2(ne141 bei2002)* | 2.85 | 1.47 | 0.74 | 0.25 |
| *mom-2(RNAi)* | 2.19 | 0.91 | 0.74 | 0.23 |
| *mom-3(or78)* | 1.09 | 0.07 | 0.12 | 0.05 |

| | | | | |
|---|---|---|---|---|
| *mom-4(ne19)* | 4.43 | 1.05 | 1.00 | 0.00 |
| *mom-4(or39)* | 3.87 | 1.02 | 1.00 | 0.00 |
| *mom-4(ne135)* | 3.13 | 0.67 | 1.00 | 0.00 |
| *mom-5(or57)* | 1.17 | 0.12 | 0.42 | 0.11 |
| *mom-5(zu193)* | 1.27 | 0.31 | 0.86 | 0.14 |
| *mom-5(RNAi)* | 1.52 | 0.48 | 0.77 | 0.16 |
| *apr-1(RNAi)* | 1.00 | 0.00 | 0.19 | 0.10 |
| *apr-1(RNAi shin1999)* | 1.00 | 0.00 | 0.15 | 0.09 |
| *dsh-2(RNAi);mig-5(RNAi)* | 1.27 | 0.31 | 0.86 | 0.14 |

Table S2: The mean and standard deviations of the parameters used in the landscape model. $P$ ratio indicates the ratio between POP-1 levels in mutant and wildtype E cells, and $S$ ratio is similarly defined.

| change from default parameters\general parameter | SKN-1 slope | SKN-1 intercept | diffusion | c_1 | c_2 | c_3 |
|---|---|---|---|---|---|---|
| 10% | 32 | 64 | 61 | 60 | 71 | 51 |
| 5% | 53 | 12 | 72 | 67 | 69 | 54 |
| 0% | 65 | 65 | 65 | 65 | 65 | 65 |
| -5% | 37 | 34 | 69 | 55 | 24 | 56 |
| -10% | 36 | 8 | 59 | 73 | 36 | 56 |

Table S3: Number of admissible wildtype POP-1, SYS-1 level solutions under perturbation of 6 general parameters.

| genotype | penetrance (%) | penetrance under 25% POP-1 reduction (%) | standard deviation (%) | penetrance under 50% POP-1 reduction (%) | standard deviation (%) |
|---|---|---|---|---|---|
| *mom-1(or10)* | 83 | 32 | 17.26 | 5 | 6.88 |
| *mom-2(or42)* | 72 | 21 | 11.26 | 3 | 4.33 |
| *mom-2(ne141)* | 53 | 11 | 5.94 | 1 | 1.58 |
| *mom-2(ne141 bei2002)* | 66 | 17 | 9.39 | 2 | 3.40 |
| *mom-2(RNAi)* | 28 | 5 | 3.99 | 0 | 0.44 |
| *mom-3(or78)* | 70 | 5 | 3.51 | 0 | 0.00 |
| *mom-4(ne19)* | 43 | 14 | 2.82 | 2 | 1.59 |
| *mom-4(or39)* | 40 | 13 | 3.42 | 1 | 1.46 |
| *mom-4(ne135)* | 11 | 3 | 1.47 | 0 | 0.06 |
| *mom-5(or57)* | 14 | 0 | 0.61 | 0 | 0.00 |
| *apr-1(RNAi)* | 29 | 1 | 0.87 | 0 | 0.00 |
| *apr-1(RNAi shin1999)* | 42 | 2 | 1.38 | 0 | 0.00 |

Table S4: Single mutant penetrance under the *pop-1* loss-of-function background as shown in Fig. 5A.

| genotype | observed penetrance (%) | penetrance from model (%) | standard deviation (%) |
|---|---|---|---|
| *sys-1(RNAi)* | 4 | 2 | 2.11 |
| *lit-1(t1534)* | 0 | 2 | 1.57 |
| *lit-1(t1512)* | 99 | 99 | 1.42 |
| *lit-1(RNAi)* | 96 | 99 | 2.06 |
| *wrm-1(RNAi)* | 100 | 99 | 1.26 |
| *mom-1(or10)* | 83 | 84 | 2.45 |
| *mom-2(or42)* | 72 | 71 | 2.45 |
| *mom-2(ne141)* | 53 | 53 | 2.54 |
| *mom-2(ne141 bei2002)* | 66 | 66 | 2.57 |
| *mom-2(RNAi)* | 28 | 27 | 2.46 |
| *mom-3(or78)* | 70 | 69 | 2.49 |
| *mom-4(ne19)* | 43 | 42 | 2.38 |
| *mom-4(or39)* | 40 | 40 | 2.50 |
| *mom-4(ne135)* | 11 | 13 | 1.95 |
| *mom-5(or57)* | 14 | 14 | 1.79 |
| *mom-5(zu193)* | 4 | 1 | 1.38 |
| *mom-5(RNAi)* | 2 | 2 | 2.06 |
| *apr-1(RNAi)* | 29 | 28 | 2.64 |
| *apr-1(RNAi shin1999)* | 42 | 41 | 2.39 |

| | | | |
|---|---|---|---|
| *dsh-2(RNAi);mig-5(RNAi)* | 4 | 1 | 1.38 |
| *mom-1(or10); sys-1(RNAi)* | 93 | 99 | 1.98 |
| *mom-2(or42); sys-1(RNAi)* | 100 | 100 | 1.33 |
| *mom-3(or78); sys-1(RNAi)* | 81 | 84 | 5.02 |
| *mom-4(ne19); sys-1(RNAi)* | 100 | 100 | 0.10 |
| *mom-5(or57); sys-1(RNAi)* | 60 | 60 | 5.44 |
| *lit-1(t1534); sys-1(RNAi)* | 100 | 98 | 2.62 |
| *wrm-1(RNAi); sys-1(RNAi)* | 100 | 100 | 0.06 |
| *mom-4(or39); mom-2(or42)* | 100 | 100 | 0.87 |
| *mom-2(ne141);mom-4(ne19)* | 100 | 98 | 2.58 |
| *mom-5(RNAi);mom-4(ne19)* | 8 | 8 | 6.10 |

Table S5: Mutant penetrance in training dataset along with the mean and standard deviation from model fitting as shown in Fig. 3A.
